# The density of doublecortin cells in the piriform cortex is affected by transition to adulthood but not first pregnancy in female mice

**DOI:** 10.1101/2023.02.08.527694

**Authors:** Rafael Esteve-Pérez, María Abellán-Álvaro, Cinta Navarro-Moreno, Michele Prina, Manuela Barneo-Muñoz, Enrique Lanuza, María José Sánchez-Catalán, Fernando Martínez-García, Jose Vicente Torres-Pérez, Carmen Agustín-Pavón

**Author notes:** Corresponding authors: JVTP and CAP. Departament de Biologia Cel·lular, Biologia Funcional i Antropologia Física, Universitat de València; C/ Dr Moliner, 50. 46100 Burjassot, València, Spain. CIBERER - IIS La Fe. These authors contributed equally to the work (Rafael Esteve-Pérez, María Abellán-Álvaro). These authors contributed equally to the work (Jose Vicente Torres-Pérez, Carmen Agustín-Pavón).

## Abstract

Motherhood is a critical period modulating behavioural changes to favour survival in mammals. In mice, olfaction is a key driver of social behaviours, and adult neurogenesis in the olfactory bulb is modulated at this stage, contributing to pup recognition. However, whether motherhood would also promote changes in the population of immature neurons of the piriform cortex is unknown. To investigate this question, we analysed the expression of doublecortin (DCX), a marker of immature neurons, in prepubescent *vs* young adults, in virgin *vs* pregnant and in pup-sensitized virgins *vs* lactating female mice. We found that the density of DCX cells sharply decreased in the piriform cortex with age, but pregnancy and lactation failed to significantly alter the density of these cells. To further analyse how motherhood could affect DCX-ir cells, we co-labelled DCX cells with NeuN, an archetypical marker of mature neurons. We did not find significant differences in the percentage of double-labelled cells nor in features related to maturation like number of neurites or main diameter in lactating dams as compared to pup-sensitized virgin females. Our results suggest that the first pregnancy does not significantly affect the differentiation of immature neurons of the piriform cortex.

## Introduction

Throughout the lifespan of a female mammal, the transition to adulthood and motherhood constitute periods of special importance due to its adaptive nature (Boivin et al. 2017; Carmona et al. 2019; Pawluski et al. 2022; Hoekzema et al. 2022). Not surprisingly, inadequate social and maternal behaviours are highly correlated with developmental and health deficits in the females and their offspring (Fitzgerald et al. 2020). Additionally, motherhood, particularly for primiparous females, represents a challenging period for the mothers themselves. Despite its importance, many studies have focused on the long-lasting consequences to the offspring while neglecting the neurobehavioral changes in the mothers (Hillerer et al. 2014; Kundakovic and Champagne 2015; Pawluski et al. 2022; Navarro-Moreno et al. 2022; Hoekzema et al. 2022) (Jodi L. Pawluski et al. 2022; Navarro-Moreno et al. 2022; Kundakovic and Champagne 2015; Hoekzema et al. 2022; Hillerer et al. 2014).

The maternal brain goes through hormonal and psychological adaptations during motherhood. The period of ‘matrescence’ will prepare the individual for the stage of motherhood, characterised by a change in behaviour (Orchard et al. 2023). For instance, female rodents start building nests and showing signs of aggression towards conspecifics by the end of their pregnancy (Mann and Svare 1982; Mayer and Rosenblatt 1984; Martín-Sánchez et al. 2015a; Navarro-Moreno et al. 2022). Similarly, the postpartum period is characterised by an increased load of caring-related tasks (e.g., nurturing and grooming the offspring) and limited or shifted interactions with other adult conspecifics (Leuner and Sabihi 2016; Pawluski et al. 2022; Navarro-Moreno et al. 2022). These behaviours are mediated by neuroplastic adaptations in the brain of the females during both pregnancy and postpartum, which are dynamically adjusted to the needs of the offspring (Fleming et al. 1994; Salais-López et al. 2020; Martínez-García et al. 2021). Thus, motherhood represents a stage of long-lasting neuroplastic adaptation in the female brain with potential recurrence during their adult life (Bridges 2016).

An evolutionary conserved architectural framework seems to orchestrate the social brain functioning in all mammals studied, the so called “socio-sexual behaviour network”, including mainly amygdaloid, septal and hypothalamic areas (Newman 1999; Swain and Ho 2019). In the maternal brain, the olfactory bulbs and hippocampus, as neurogenic areas, have also been shown to play a relevant role (Leuner and Sabihi 2016). Adult neurogenesis, the production of new neurons from adult neural precursors, has emerged as key contributor to parental neuroplasticity in this circuitry. This process occurs mainly in the sub-granular zone (SGZ) of the hippocampal dentate gyrus (DG) and the ventricular-subventricular zone (V-SVZ), which gives rise to the neuroblasts that migrate via the rostral migratory stream (RMS) towards the olfactory bulb, where they disperse and integrate as inhibitory interneurons (Doetsch and Alvarez-Buylla 1996; Leuner and Sabihi 2016). Adult neurogenesis in these niches is sensitive to hormonal changes occurring during pregnancy and lactation and has been involved in the expression of maternal behaviours (Shingo et al. 2003; Furuta and Bridges 2005; Pawluski et al. 2009; Mak and Weiss 2010; Leuner and Sabihi 2016; Medina and Workman 2020). Interestingly, a recent study showed that pregnancy stimulates adult neurogenesis in the olfactory bulb, contributing to pup recognition (Chaker et al. 2023).

Studies in different species indicate that doublecortin (DCX), a protein expressed by immature neurons, can also be observed in non-proliferative areas of the olfactory system in the adult brain. Immature neurons of embryonic origin are located in layer II of the piriform cortex (Pir) (Nacher et al. 2002; Rubio et al. 2016; La Rosa et al. 2020), and remain in a posed neuroblast-like state (arrested maturation), slowly maturing with age (Ghibaudi et al. 2023b), and integrating as glutamatergic neurons (Rotheneichner et al. 2018). Some studies have shown that the maturation rate of DCX cells can be affected by olfactory deprivation (Gómez-Climent et al. 2011), stress (Nacher et al. 2004; Abellán-Álvaro et al. 2024) or pharmacological manipulations (Coviello et al. 2020). However, whether motherhood affects this population is unknown.

Thus, here we sought to investigate the potential influence of the first pregnancy on DCX-ir cell density in the Pir. We first assessed the effect of transition to adulthood by comparing prepubescent *vs* young adult female mice with full reproductive capacity (Arellano et al. 2024). Secondly, we investigated the effect of pregnancy by comparing young adult virgin *vs* pregnant mice, and the effect of lactation by comparing pup-sensitized virgins vs lactating dams (Martín-Sánchez et al. 2015b). We hypothesised that both transition to adulthood and motherhood would promote maturation, which could be translated as a decrease on the number of immature embryonically-generated DCX-ir neurons at Pir.

## Materials and Methods

### Animals

We used 42 female mice (*Mus musculus*) from the CD1 strain. Their age at the time of sacrifice was 4 weeks old for the prepubescent animals, and 12-13 weeks old for the young adult virgin, pregnant, pup-sensitized virgin and dam groups. Subjects were housed in age-matched homologous pairs to avoid isolation stress and kept in polypropylene cages at ∼24°C under a 12-h light/dark hour cycle (lights on at 8:00 am) with *ad libitum* access to water and food.

Animal procedures were approved by the Committee of Ethics and Animal Experimentation of the Universitat Jaume I, performed in accordance with the European Union Council Directive of June 3^rd^, 2010 (6106/1/10 REV1), and under an animal-usage license issued by the Direcció General de Producció Agrària i Ramaderia de la Generalitat Valenciana (code 2015/VSC/PEAI00055 type 2).

### Experimental design

Subjects were randomly assigned to the different experimental groups. For Experiment 1, we used prepubescent (n = 7) *vs* young adult virgin females (n = 7). For Experiment 2, we analysed used virgin (n = 8) *vs* pregnant (gestational day 17, n = 7) females. This samples were parallel series of the samples used elsewhere (Navarro-Moreno et al. 2020). Finally, for Experiment 3 we used pup-sensitised virgins (n = 6) *vs* lactating dams (postpartum day 4, n = 7). For this experiment, females of 10 weeks old were paired with a stud male for three days. When pregnancy was confirmed, each pregnant female was housed with an age-matched virgin female, which remained with the dam since then until the completion of the study at 13 weeks of age. This procedure allows the virgin females to be pup-sensitized, thus controlling that any changes seen in dams were due to motherhood and not to mere contact with pups (Martín-Sánchez et al. 2015b).

In all cases, at corresponding age, females were deeply anaesthetised with an intraperitoneal (i.p.) injection of sodium pentobarbital (Vetoquinol, Madrid, Spain) at 0.02 mg per g of body weight, and transcardially perfused with saline (0.9%) 4% paraformaldehyde (PFA) in 0.1 M phosphate buffer (PB), pH 7.4.

### Tissue processing

After perfusion, brains were carefully extracted from the skull, post-fixed overnight at 4°C with similar fixative solution and kept in a cryopreserving solution of 30% sucrose in 0.01 M PB at 4°C until sinking. Five parallel series of 40 μm-thick free-floating coronal sections were obtained from each brain using a freezing microtome (Microm HM-450, Walldorf, Germany). Sections were kept in 30% sucrose in 0.1 M, pH 7.6, at −20°C until use.

### Immunofluorescence

We used one out of the five brain series from each subject. Sections corresponding to each Experiment were simultaneously processed using same batches of reagents and antibodies to ensure minimal procedural biases. Immunofluorescence was performed in free-floating conditions. First, endogenous fluorescence was blocked by incubating the sections in 1% NaBH in 0.05 M tris-buffered saline (TBS) for 30 minutes at room temperature (RT). Next, to block nonspecific binding, sections were incubated in a 3% mixture of normal donkey serum (NDS) and normal goat serum (NGS) in 0.3% Tx100 in 0.05 M TBS, pH 7.6, for one hour at RT. Then, sections were incubated overnight with the primary antibody against DCX (Guinea pig anti-DCX, 1:4000; EMD Millipore Corp., AB2253), or a mix of DCX and NeuN (Mouse anti-NeuN, 1:2500; EMD Millipore Corp., MAB377), diluted in 0.05 M TBS, pH 7.6, with 2% NDS/NGS in 0.3% Tx100 at 4°C in agitation. Following day, sections were incubated with the secondary antibodies (1:200 each; Alexa Fluor 488-conjugated goat anti-guinea pig, Molecular Probes, for DCX; Rhodamine Red^TM^-X-conjugated donkey anti-rabbit, Jackson ImmunoResearch, for NeuN) diluted in 0.05 M TBS, pH 7.6, with 2% NDS in 0.3% Tx100 for 90 minutes at RT. When required, samples were thoroughly rinsed between steps in TBS.

To fluorescently label nuclei, sections were also incubated with 600 nM DAPI (4’,6’-diamino-2-feniindol) in TBS for 45 seconds at RT. Finally, sections were mounted in glass slides using 0.2% gelatine in TB and cover-slipped with fluorescent mounting medium (FluorSave Tm Reagent; Dako, Glostrup, Denmark).

### Image acquisition and quantification

In all Experiments, we quantified the number of DCX-ir cells in 3-4 representative sections of the Piriform cortex (Bregma +1 mm, 0 mm, -1 and -2 mm). Pictures were obtained by an experimenter blind to the experimental groups from both hemispheres at selected levels centred at layer II. All images were taken using digital DFC495 camera attached to a Leitz DMRB microscope (Leica, Wezlar, Germany). Unless stated otherwise, pictures were taken at 20x magnification. Positive DCX-ir cells at Pir (layer II, occasionally migrating towards layer III) were counted by using the multi-point tool of ImageJ analysis software. Results were averaged and expressed as number of positive cells per mm^2^. Besides, the main diameter of these DCX-ir cells in the layer II of the Pir was estimated for a posterior classification between tangled (main diameter <11µm, more immature) or complex (main diameter >11µm, in a more advanced maturation stage), similarly to what has been done in other studies (Coviello et al. 2021).

We further analysed DCX-NeuN immunofluorescence in Experiment 3. To study co-localization, representative images of the Pir at the selected levels were taken with a confocal microscope (ZEISS LCS 980) at 20x magnification. Quantification of the percentage of DCX-ir that co-expressed NeuN-ir was conducted manually throughout the whole II layer of the Pir available in each image. Similarly, the main diameter of each DCX-ir cell in layer II, as well as their number of neurites, were determined manually. Figures were processed using Fiji software (NIH) and Adobe Photoshop CS6 (Adobe Systems, MountainView, CA, USA)

### Statistical analysis

Data were analysed using the software IBM SPSS Statistics 26.0. Normality was determined using the Shapiro-Wilk tests. We applied a student’s t-test analysis for independent samples when possible, and if data violated normality, we used a Mann-Whitney test. Significance was set al p < 0.05. Graphs were created using ggplot2 package (Wickham 2016) in R (version 4.2.1; (Team 2023).

## Results

As expected, we found DCX-ir labelled cells in layer II of the Pir in all the analysed samples. These cells could be identified as tangled (small somata and one or two neurites) or complex, in a more advanced phase of maturation (Figure 1A). In Experiment 1, comparing prepubescent females with young adult females, we found a sharp and significant decrease in the density of DCX-ir cells (W = 45.5, p = 0.0086; Figure 1B). However, the percentage of complex cells (main diameter >11µm) DCX-ir cells did not significantly differ with age, suggesting a uniform maturation process (W = 21; p = 0.7; Figure 1A; Figure 1C).

**Figure 1.**
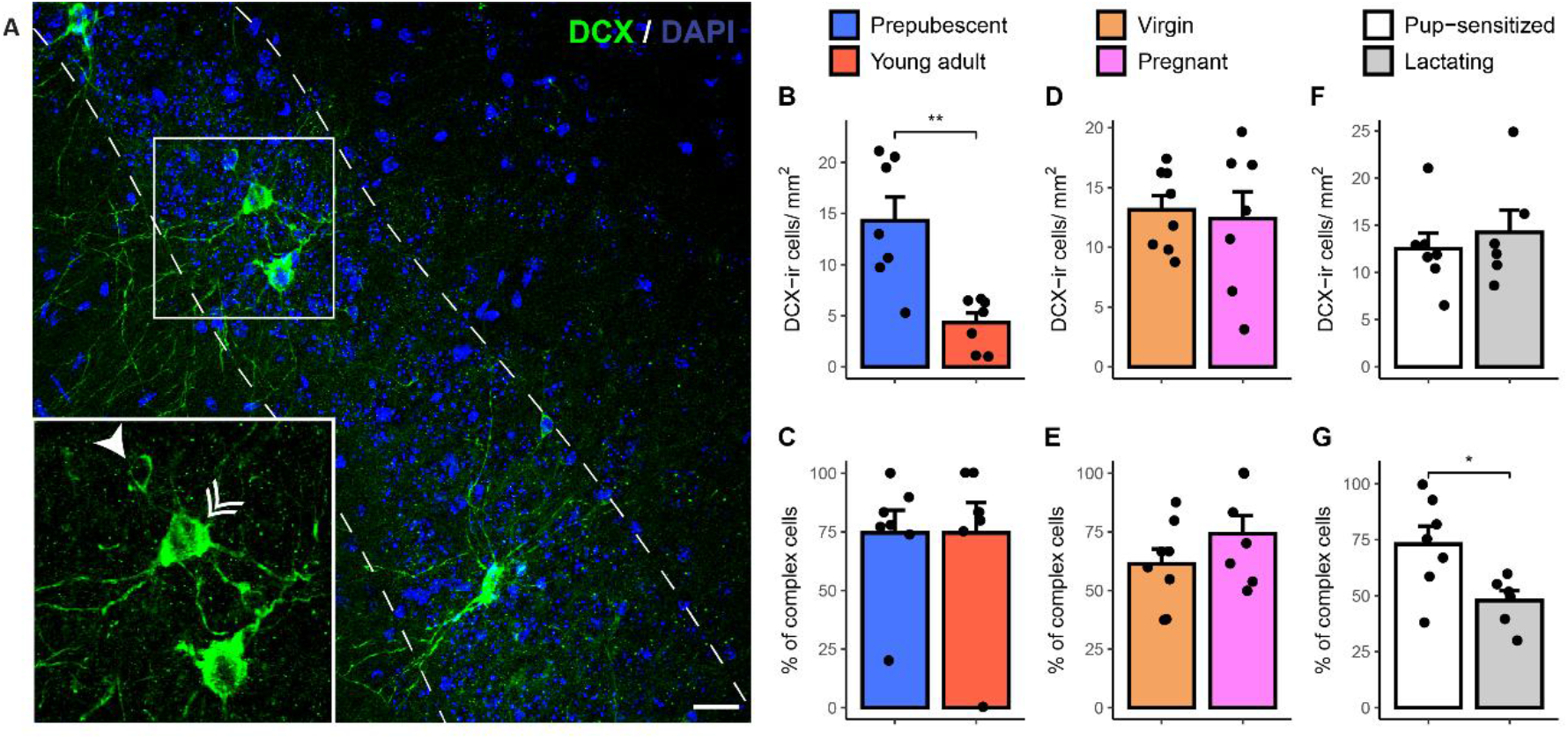
DCX-ir cells at the piriform cortex (Pir) throughout age and motherhood. **A)** Representative images, and squared magnification, of DCX (green) and DAPI (blue) immunoreactivity in the Pir, with tangled (single arrow) and complex (double arrow) DCX-ir cells in Pir’s layer II (dashed lines). Scale bar: 20 µm. Effect of **B)** transition to adulthood (prepubescent -blue-vs. young adult -red), **D)** pregnancy (virgin -orange-vs. pregnant -pink), and **F)** lactation (pup-sensitized – white-vs. lactating -grey) in the density of DCX-ir cells at Pir. Effect of **C)** puberty, **E)** pregnancy, and **G)** lactation in the percentage of complex DCX-ir cells at Pir. All graphs represent mean + Standard Error Mean (SEM), and individual values (dots). **: p< 0.01.

By contrast, in Experiment 2, we did not find any significant difference when comparing virgin and pregnant mice in the density of DCX-ir cells in the layer II of the Pir (t = 0.3, p = 0.77; Figure 1D) nor in the percentage of complex DCX-ir cells (t = -1.28, p = 0.22; Figure 1E). Similarly, in Experiment 3 we could not find any significant change in density (t = 0.62, p = 0.28; Figure 1F), but we found a slight decrease in the percentage of complex DCX-ir cells (t = -2.63, p = 0.012; Figure 1G) in lactating mice as compared to pup-sensitized virgins.

Since the decrease in the percentage of complex cells could point towards an accelerated disappearance of DCX in complex cells closer to maturation, we next explored the co-labelling of DCX with NeuN. As expected, we found that some of the DCX-ir neurons co-expressed NeuN, suggesting their transition to mature neurons (Figure 2A). However, the percentage of these double-labelled cells was not significantly different between virgin and lactating mice (t = 1.01, p = 0.34; Figure 2B). We also assessed other features related to maturation, such as the number of neurites and the size of the soma, but we did not find significant differences between groups on those measures (number of neurites, t = 1.41, p = 0.19; Figure 2C, main diameter t = 1.52, p = 0.16; Figure 2D).

**Figure 2.**
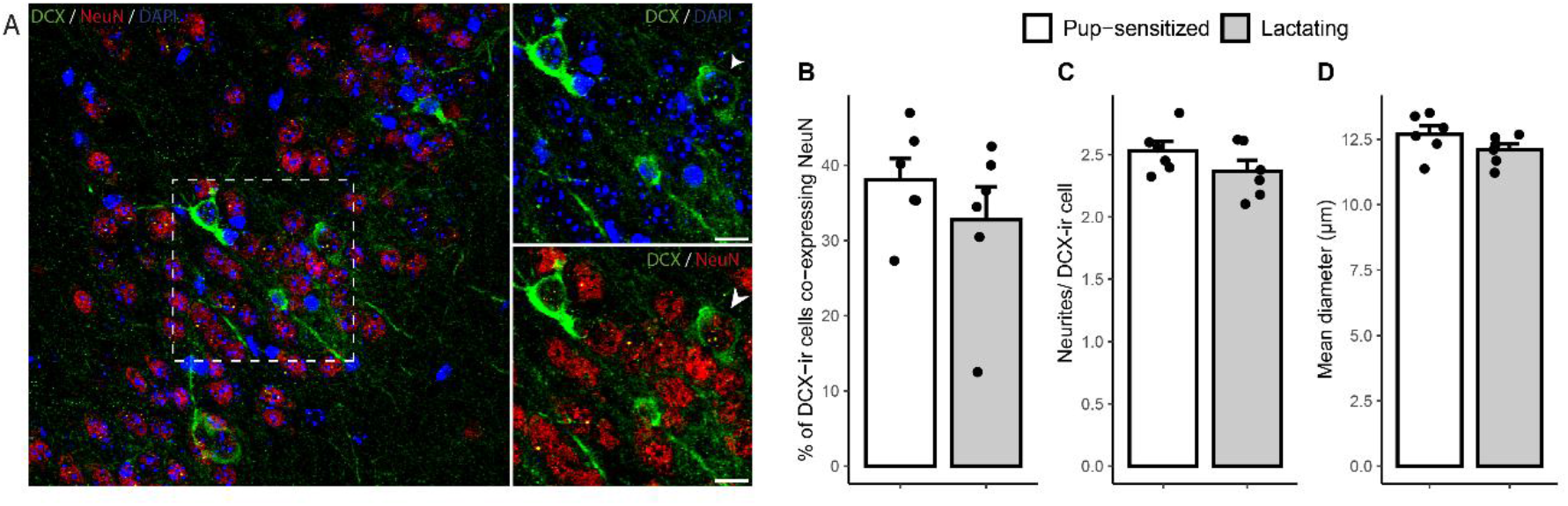
DCX-ir co-expression with NeuN is not affected by lactation in the piriform cortex (Pir). **A)** Representative images of DCX (green), NeuN (red) and DAPI (blue) immunoreactivity in the layer II of the Pir, with an example of double-labelled cell (arrow). Dashed square (left) indicates magnified area (right). Scale bar: 10 µm. Effect of lactation (pup-sensitized in white vs. lactating in grey) on the percentage of **B)** double-stained cells, **C)** number of neurites per DCX-ir cell, and **D)** main diameter of DCX-ir cells. All graphs represent mean + Standard Error Mean (SEM), and individual values (dots).

## Discussion

Our study is in agreement with previous findings showing that age leads to a decrease in the density of DCX-ir cells in the Pir, that lose immaturity markers as they differentiate into glutamatergic neurons. However, we could not find a significant effect of the first pregnancy in the density of these cells.

Motherhood has been previously associated with neuroplastic changes including adult neurogenesis at the V-SVZ which, via the canonical pathway, migrate to the OB and integrate as mature neurons (Shingo et al. 2003; Leuner and Sabihi 2016; Medina and Workman 2020). These plastic changes are adaptive since they are associated with novel odour learning, including pup recognition and maternal care, hence increasing the potential of the offspring to survive (Hillerer et al. 2014; Chaker et al. 2023). Given the olfactory and integrative functions of the Pir, as well as its connections and its reported adaptive capacities (Chen et al. 2022), we expected that a considerably strong adaptive process like ‘matrescence’ would also result in an increased maturation and integration of this population of cells in the Pir. However, in disagreement with our hypothesis, we did not find a significant impact of pregnancy or lactation on the density of these cells, and just a slight change in the percentage of complex DCX-ir cells in lactating females, suggesting a minor loss of complex cells.

The population of DCX-ir cells at the Pir has been previously described as a niche for protracted neural maturation, where DCX-ir cells decrease with age as they mature and integrate into the neural circuit (Gómez-Climent et al. 2008; Rubio et al. 2016; Rotheneichner et al. 2018; La Rosa et al. 2020; Ghibaudi et al. 2023b). In our study, we found a sharp decline in the density of DCX-ir cells in young adult virgins (roughly 3 months old) as compared to prepubescent females (1 month old), suggesting an important effect of pubertal development in the maturation rate of DCX-ir cells. Albeit following the same trend, a previous study failed to demonstrate a significant decrease in the density of DCX-ir cells until older adulthood, i.e, when comparing 1 month old *vs* 12 months old mice (Ghibaudi et al. 2023b). However, that study was performed in C57/BL6 male mice, so factors such as the sex of the animals, the strain and the method of quantification might account for the observed differences in the time course of maturation.

More surprising to us was the absence of statistically significant motherhood-induced differences in DCX expression in the Pir. We hypothesise that these females being primiparous might be an important factor leading to lack of a significant effect. In the DG of the hippocampus, the pattern of cell proliferation depends on both the reproductive state and the reproductive experience and slightly changes depending on whether the mother is primiparous or multiparous (Pawluski and Galea 2007). Therefore, in the Pir, repetition of ‘matrescence’ cycles might result in a stronger, and quantifiable, effect. In addition, the pattern of DCX-ir cell maturation in the Pir is different than that of the V-SVZ or the DG of the hippocampus, with their DCX-ir immature neurons having an embryonic origin and lacking proliferative capacity (Gómez-Climent et al. 2008; Rubio et al. 2016; Rotheneichner et al. 2018). These different patterns of generation and/or maturation might explain why we have not seen an apparent effect of pregnancy and lactation in the density of DCX-ir cells. Alternatively, it is possible that the time at sacrifice, postpartum day 4, might be too early to see significant differences in the density of a population not affected during pregnancy. In this sense, we detected a slight effect on the composition of DCX-ir population, namely a decrease in the percentage of complex DCX-ir cells, the ones in a more advanced status of maturation. Of note, pregnancy-induced increase in neurogenesis in the OB is transient, with new-born granular cells disappearing by weaning of the litters (Chaker et al. 2023). It is possible then that only a small amount of complex DCX cells might be recruited on the first pregnancy, hence masking any significant effect on the average density of the whole population.

We have identified some potential caveats. Firstly, despite being considered a microtubule-associated protein with functions on differentiation and migration of neurons (Francis et al. 1999; Gleeson et al. 1999; Friocourt 2003), DCX could be involved in other processes which might be related to neuronal development (Yap et al. 2012) or not (Dhaliwal et al. 2016). Thus, interpretations on neuronal maturation based on DCX-ir alone should be done with caution. Secondly, different commercial antibodies against DCX might yield variations on data depending on the species and cerebral area studied (Ghibaudi et al. 2023a), a source of inconsistency that should also be accounted for. In addition, the findings reported here are just descriptive. Future studies using viral labelling or genetically modified mice which will allow for traceability of DCX neurons, multiple pregnancies and later time points of the lactating process are needed to understand the dynamics of maturation, integration and its functional significance.

Many studies have focused on the effect of motherhood to the offspring while failing to look after the consequences to the mothers themselves. Different imaging techniques have allowed identification of structural and functional neuroplastic adaptations associated with motherhood in humans (Hoekzema et al. 2017; Servin-Barthet et al. 2025). However, these studies can only provide limited data and do not allow for cellular-type identification. Here, we have used a rodent model to study how the population of immature neurons of the Pir is affected by motherhood. Apart from replicating and extending previous findings showing a sharp decrease of DCX-ir neurons in the Pir after puberty, we have found that pregnancy and lactation do not significantly alter the maturation and/or integration of these DCX-ir immature neurons, maybe due to the number of reproductive experiences and/or Pir’s peculiarities in terms of maturation. Therefore, additional maturation inputs, potentially related to olfaction, should be studied to decipher how and why this population, which remains in an immature state, starts to mature.

## Data availability

Raw data generated within this manuscript can be requested to the corresponding authors.

## Author contributions

RE-P, and MA-A performed the immunofluorescence procedures, acquired images, analysed data, figure preparation and contributed to writing. CN-M and MB-M performed animal experiments and tissue processing. MP acquired images and analysed data. MJS-C, FM-G and EL participated in the study design and contributed to writing and funding acquisition. JVT-P participated in image acquisition, data analysis, prepared figures, and wrote the manuscript. CA-P obtained funding, designed the study, supervised research, performed data analysis and wrote the manuscript. Final version of this manuscript was discussed and approved by all authors.

## Acknowledgments

We thank Dr Silvia De Marchis (Università degli Studi di Torino) and Dr Vicente Herranz-Pérez (Universitat de València) for critically reviewing a previous and latest version of this manuscript respectively, and Mr Josep Pardo García and Mr Samuel Martín for technical assistance.

